# Electrical Synapse Rectification Enables Dual-Network Activity in the crab *Cancer borealis*

**DOI:** 10.1101/2024.08.15.608183

**Authors:** Savanna-Rae H Fahoum, Farzan Nadim, Dawn M Blitz

**Affiliations:** Department of Biology Center for Neuroscience and Behavior Miami University Oxford, OH 45056; Federated Department of Biological Sciences New Jersey Institute of Technology and Rutgers University Newark, NJ 07102

**Keywords:** Central Pattern Generator, Neuropeptide, Neuronal Switching, Oscillations, Synchrony

## Abstract

Flexibility of rhythmic networks includes neuromodulator-elicited changes in neuronal participation between networks. We are examining the role of rectifying electrical synapses in this neuronal switching. Electrical synapses can have complex, non-intuitive effects on network output. However, it is often difficult to measure and manipulate rectification across conditions to determine their functional contributions. Here, we use the Jonah crab *Cancer borealis* to investigate rectification in well-described rhythmic networks. In an established modulatory state, stimulating the projection neuron MCN5 or bath applying its neuropeptide Gly^1^-SIFamide causes the two LPG neurons to switch from pyloric rhythm-only (food filtering, ∼1 Hz) activity to dual pyloric and gastric mill rhythm (chewing, ∼0.1 Hz) activity. Typically, LPG is co-active with the two PD and single AB pyloric pacemaker neurons due to rectifying electrical coupling. In Gly^1^-SIFamide, LPG continues to burst in pyloric time with AB/PD but periodically “escapes” and generates intrinsic longer-duration gastric mill-timed bursts, decreasing its overall synchrony with AB/PD, while AB/PD retain their synchronous pyloric timing. Using two-electrode voltage clamp recordings, we find that Gly^1^-SIFamide does not alter electrical coupling strength or the rectification between LPG and AB/PD. However, in a computational model, rectification is necessary for LPG to escape AB/PD electrical coupling and generate longer, gastric mill-timed bursts. This was confirmed in the biological system by adding a dynamic clamp non-rectifying electrical synapse between LPG and PD, which decreased LPG’s escape from AB/PD and its gastric mill-timed activity. Thus, rectification between electrically coupled oscillators can underlie modulator-elicited changes in their synchrony.

**Significance statement:** Behavioral adaptability includes rapid changes in neural circuit composition. Unlike chemical synapses, little is known about electrical synapse contributions to circuit flexibility. Here, we identify an important role for electrical coupling rectification (asymmetrical current flow between coupled neurons) in regulating neuronal participation between neural circuits. A neuromodulator causes the activity of two sets of electrically-coupled neurons participating in an oscillatory neural circuit to diverge, allowing one set to additionally participate in a second oscillatory circuit at a distinct frequency. Using mathematical modeling and the dynamic clamp technique, we show that this transition depends on rectification. This newly-identified function of electrical synapse rectification may have implications for oscillatory networks underlying a range of functions, due to the prevalence of electrical coupling.

## Introduction

Oscillatory neural networks underlying rhythmic motor behaviors, or sensory or higher order neural processing are flexible to enable organisms to adapt to changes in their physiological state and external environment (1–4). This network plasticity includes neuronal switching, in which neuromodulatory inputs can alter the participation of neurons, effectively “moving” neurons from one network to another, or between single- and dual-network participation. Such changes in the functional composition of networks occurs due to modulation of chemical synapses or intrinsic properties (5–9). In addition to chemical synapses, electrical synapses are prevalent components of neural networks (10–12). However, little is understood about the contribution of rectifying electrical synapses to network function (10, 13–16).

Electrical synapses exhibit rectification when current flows better in one direction than the other. Rectification may be complete or partial, and the extent of rectification is subject to change (17–19). Rectification may occur due to structural differences such as different hemichannel proteins (19, 22, 23), or differences in the intrinsic properties, ion concentrations, or synaptic support proteins of the connected neurons (17, 19). Originally identified in the crayfish escape system, rectification is now known to occur in many animals and many brain regions (17–21). Nevertheless, the prevalence of rectification is still likely underestimated because of the difficulty of dual recording from coupled neurons and, as computational studies indicate, with somatic recordings, synapse location and intrinsic neuronal properties can mask rectification (17, 21). Consequently, the functional role of rectifying electrical synapses and their role in circuit flexibility is poorly understood. In general, electrical synapses are commonly thought to contribute to synchronization, even though they can, unintuitively, also contribute to antiphase activity of neurons in rhythmic networks or even disruption of oscillations (14, 16, 17, 24–26). Strongly rectifying electrical synapses can play a role when rapid unidirectional signaling is important, such as in auditory processing or escape circuits (17, 19, 27). Additionally, computational studies suggest interesting, complex contributions of rectifying electrical synapses to network dynamics, such as regulating the sensitivity of a network to the strength of its chemical synapses (15).

Here we take advantage of small well-defined neural circuits, and an identified modulatory projection neuron with identified cotransmitters, to investigate whether rectifying electrical synapses enable neurons from one circuit to participate in the activity of a second distinct circuit (8, 28, 29). The crab stomatogastric ganglion (STG) includes two rhythmic networks with very different cycle frequencies: the slower gastric mill (∼0.1 Hz) and the faster pyloric (∼1 Hz) networks (2, 30). Some STG neurons are dedicated participants in only one network, while others may participate in both or switch their participation based on the modulatory state (31, 32). For instance, appropriate modulatory inputs enable the lateral posterior gastric (LPG) neuron to switch from its control state of only fast pyloric oscillations to dual slow/fast (gastropyloric) oscillations (8, 29, 33). We have previously shown that the slower gastric mill-timed bursts depend on the modulation of LPG intrinsic properties by the neuropeptide Gly^1^-SIFamide (8, 28), but it is unclear how these long bursts supplant the fast pyloric-timed activity, while its electrically-coupled partner neurons remain pyloric timed. We hypothesized that LPG’s ability to produce the slower bursts requires 1) decreased strength of electrical coupling, and 2) rectification of the remaining electrical coupling.

## Results

### Pyloric and Gastric Mill Networks

The pyloric network is pacemaker-driven and continuously active in vivo, or in vitro, unless neuromodulatory inputs are removed (2, 30). The pyloric pacemaker ensemble includes the single anterior burster (AB) neuron, two pyloric dilator (PD) neurons, and two LPG neurons (Fig. 1B, *left*) which produce synchronous bursting oscillations due to strong electrical coupling (Fig. 1A-B) (14, 34, 35). The electrical synapses between AB:LPG and PD:LPG are rectifying, with greater positive current flowing from AB and PD to LPG. Within a neuron type (PD:PD and LPG:LPG) the coupling is symmetric (35, 36). Finally, there may be a weak rectification of the AB:PD synapse (35) but as yet, this connection is not fully examined.

**Figure 1.**
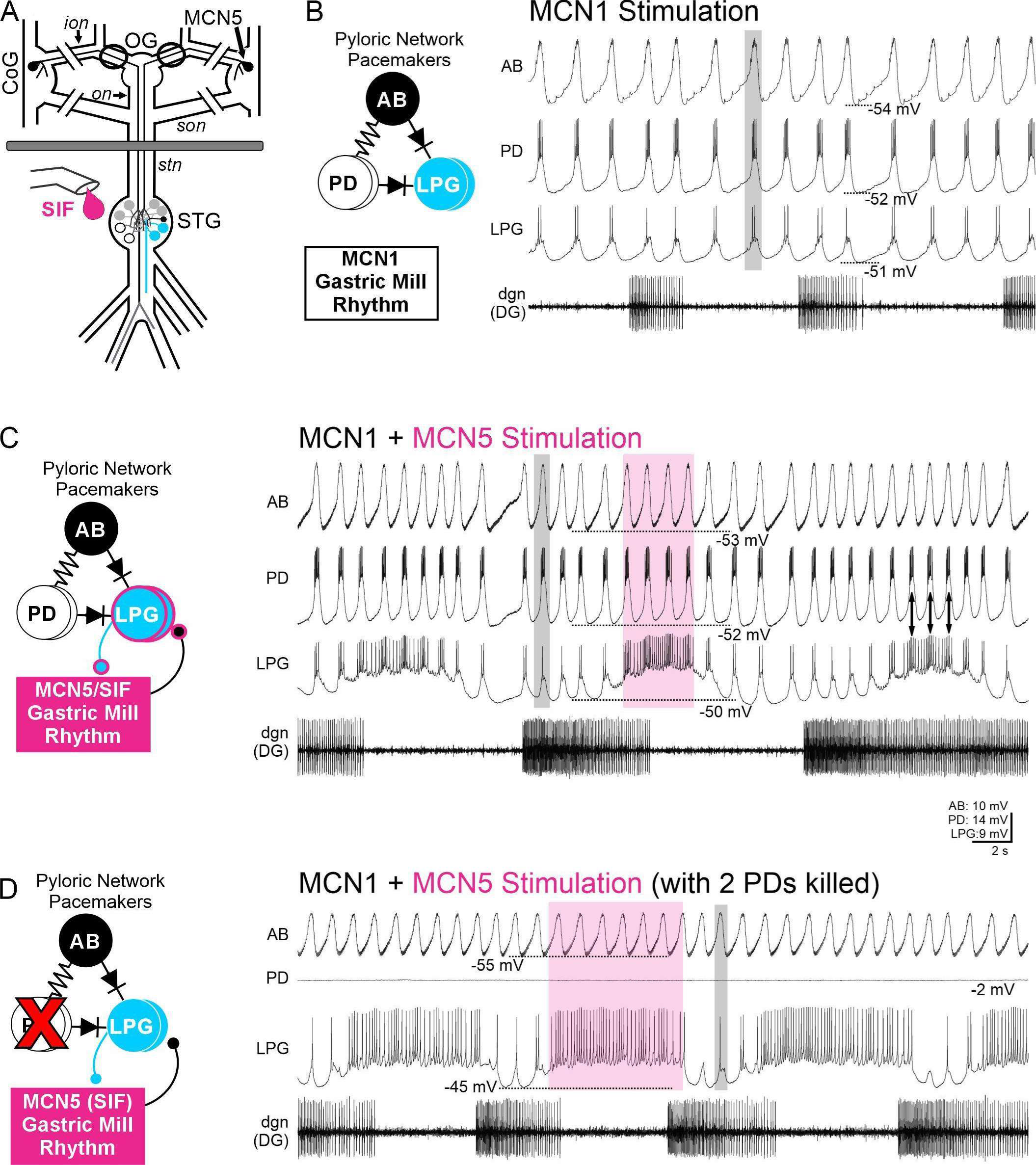
The modulatory neuron MCN5 or bath application of its neuropeptide transmitter Gly^1^-SIFamide enables the LPG neuron to periodically escape from the rest of the pyloric pacemaker neurons, including escaping from direct coupling to the AB pacemaker neuron. A) The schematic of the isolated stomatogastric nervous system includes the paired commissural ganglia (CoGs) and single oesophageal ganglion (OG), sources of descending modulatory inputs, the stomatogastric ganglion (STG) containing CPG circuits for the pyloric and gastric mill rhythms, and their connecting and peripheral nerves. B) *Left*, The circuit diagram indicates the connectivity among the pyloric pacemaker ensemble neurons including the pacemaker interneuron AB (1 copy), PD motor neurons (2 copies), and LPG motor neurons (2 copies). These neurons are connected through electrical synapses, including rectification (diode symbols) of the LPG connections to AB and PD, which are the focus of this study. In the control condition, AB, PD, and LPG are active in pyloric time due to their electrical coupling, and there are no functional synaptic connections between the gastric mill network and the LPG neuron. *Right*, MCN1 stimulation (15 Hz) was used to activate a pyloric rhythm as a baseline to which the MCN5/SIF state could be compared in this example. Intracellular recordings of AB, PD, and LPG monitor the pyloric rhythm. All three neurons were oscillating synchronously (example simultaneous depolarizations highlighted with gray box). The MCN1-elicited gastric mill rhythm does not include recruitment of LPG into the gastric mill network. C) *Left,* MCN1 and MCN5 co-stimulation (30 Hz) elicits MCN5/Gly^1^-SIFamide (MCN5/SIF) modulatory actions including activation of intrinsic bursting in the LPG neuron (pink circle around LPG neurons) and enhancement of bidirectional inhibitory chemical synapses between LPG and gastric mill neurons (lines and balls with pink outlines in schematic). *Right*, During MCN1 and MCN5 co-stimulation, MCN5/SIF modulation enables dual-network activity in which LPG continues to oscillate in time with AB and PD (e.g., gray box) but periodically escapes and generates longer duration, gastric mill-timed bursts that persist for multiple AB/PD oscillations (e.g., pink box). D) *Left*, The two PD neurons were photoinactivated (X symbol; see Methods) to test whether AB and LPG are directly coupled, or only coupled through the PD neurons, and whether LPG was able to generate dual-network activity when only coupled to the AB neuron. *Right*, The PD neuron recording shown, along with the second PD neuron were photoinactivated, evident by the membrane potential reaching approximately 0 mV (see Methods). During MCN1 and MCN5 co-stimulation, AB and LPG appear to be electrically coupled due to their synchronous pyloric-timed activity (grey box). The only other pyloric neuron known to influence LPG is the LP neuron, which was photoinactivated in this experiment. LPG also generated gastric mill-timed bursting (pink box) with both PD neurons photoinactivated. Recordings in B-D are from the same experiment. Scale bars apply to all panels.

To establish a controlled baseline condition, we removed descending modulatory inputs by transecting the nerves through which they project (*ion*s and *son*s, Fig. 1A). In this condition, the pyloric rhythm slows or shuts off (37, 38). To reactivate the pyloric rhythm for an internal control to which we could compare the modulated pacemaker neuron activity, in some preparations we stimulated the modulatory commissural neuron 1 (MCN1) (15 Hz) to reactivate the pyloric rhythm, during which the AB, PD, and LPG neurons recovered their synchronous activity (Fig. 1B, one set of bursts highlighted with grey box).

In contrast to the pyloric rhythm, the gastric mill rhythm is not driven by a pacemaker, but is generated by a half-center oscillatory pair of neurons (2, 30). The gastric mill rhythm is episodic, requiring a stimulus for activation and, in the crab *Cancer borealis*, multiple versions of the gastric mill rhythm can be elicited by sensory input or stimulation of identified modulatory neurons (42–44). For example, activation of the modulatory commissural neuron 5 (MCN5) or bath-application of its neuropeptide transmitter Gly^1^-SIFamide (SIF) elicits a unique gastric mill rhythm (8, 29, 33, 45) which we will refer to as the MCN5/SIF modulatory state. Here we focus on one aspect of this modulatory state, the switching of the LPG neuron from pyloric time-only activity, to dual pyloric and gastric mill-timed activity (Fig. 1B-D) (8, 28, 29, 33).

### A switch from single- to dual-network activity

MCN5/SIF modulates ionic currents (Fig. 1C, pink circles around LPG) to enable LPG to produce slower, longer duration intrinsic bursts, which are coordinated into a gastric mill rhythm due to parallel Gly^1^-SIFamide modulation of bidirectional chemical synapses between LPG and gastric mill neurons (Fig. 1C, pink circles around synapses) (8, 28, 46). In the MCN5/SIF modulatory state, LPG continues to produce pyloric bursts in time with AB and PD (Fig. 1C, *right* grey box) but periodically escapes its pyloric pacemaker-timed activity to produce longer-duration gastric mill-timed bursts (Fig. 1C, *right* pink box). During LPG gastric mill bursts, AB and PD continue to oscillate in pyloric time. When there is an LPG escape, the AB and PD trough voltages remain mostly the same (Fig. 1C, AB, PD, dotted lines), whereas the LPG trough voltage is depolarized throughout each gastric mill burst, compared to during its pyloric-timed activity (Fig. 1C, LPG, dotted line). Although their voltage waveforms diverge during LPG gastric mill bursts, it appears that LPG remains coupled to AB/PD, as the frequency of the AB/PD oscillations and thus the pyloric rhythm is faster during these long LPG bursts (Fig. 1C) (45). Maintained coupling is also suggested by the periodicity of the LPG action potential frequency in time with AB/PD depolarizations during the LPG gastric mill bursts (Fig. 1C, double arrows). We will address the coupling strength among the pyloric pacemaker neurons during the MCN5/SIF modulatory state below.

While each of the five pyloric pacemaker ensemble neurons is influenced by the other four (35), it was not known if LPG is directly coupled to the AB pacemaker neuron, or if its coupling occurs via the PD neurons. To test for direct electrical coupling between AB and the LPG neurons, we photoinactivated (see *Methods*) the two PD neurons and recorded from AB and LPG. We found that AB and LPG were co-active in the absence of the PD neurons in control conditions (n=14/14). Within this dataset, AB and LPG were synchronously active when the LP neuron, which provides the only (inhibitory) feedback synapse onto the pacemaker neurons in control conditions, was photoinactivated (n=4) or hyperpolarized (n=2), or when inhibition was blocked with picrotoxin (PTX, 10 µM; n=2). Thus, LPG appears to be directly electrically coupled to the pyloric pacemaker neuron, AB. We also tested whether LPG and PD were directly coupled: after AB was photoinactivated, depolarizing or hyperpolarizing current injections in PD or LPG resulted in smaller amplitude depolarizations or hyperpolarizations, respectively, in the other neuron (n=3/3; *SI Appendix*, Fig. S1). Therefore, LPG appears to be directly electrically coupled to both AB and PD, and therefore must escape this coupling to generate the longer-duration gastric mill bursts in the MCN5/SIF state.

To quantify the functional coupling of LPG, AB and PD neurons, we measured the pairwise correlation coefficient of their voltage waveforms (47, 48) over 100-200 s time blocks in control (saline or MCN1 stimulation) vs MCN5/SIF (Gly^1^-SIFamide bath application or MCN1+ MCN5 stimulation). In an example preparation, PD and LPG generated simultaneous pyloric-timed bursts in saline, but produced different numbers of action potentials (Fig. 2Ai). A plot of LPG voltage against PD voltage indicated correlated slow wave oscillations (∼-55 to -40 mV), but less correlation of their individual action potentials, evident in the divergence of their waveforms at more depolarized voltages (Fig. 2Aii). In the MCN5/SIF state when LPG was generating gastric mill bursts, the pairwise correlation of the PD:LPG voltage waveforms was weaker (Fig. 2Aiii, Aiv). Across preparations, synchrony consistently decreased between LPG:PD and between LPG:AB from saline/MCN1 to MCN5/SIF (Fig. 2C) (LPG:PD, t(9)=6.29, p=0.00014, paired t-test, n=10; LPG:AB, t(3)=5.78, p=0.010, paired t-test, n=4). In contrast, there was no change in the synchrony of AB:PD (example experiment: Fig. 2Bi-iv) or PD:PD pairs between saline and Gly^1^-SIFamide (Fig. 2C) (AB:PD, t(3)=1.33, p=0.28, n=4; PD:PD, t(2)=0.71, p=0.55, n=3).

**Figure 2.**
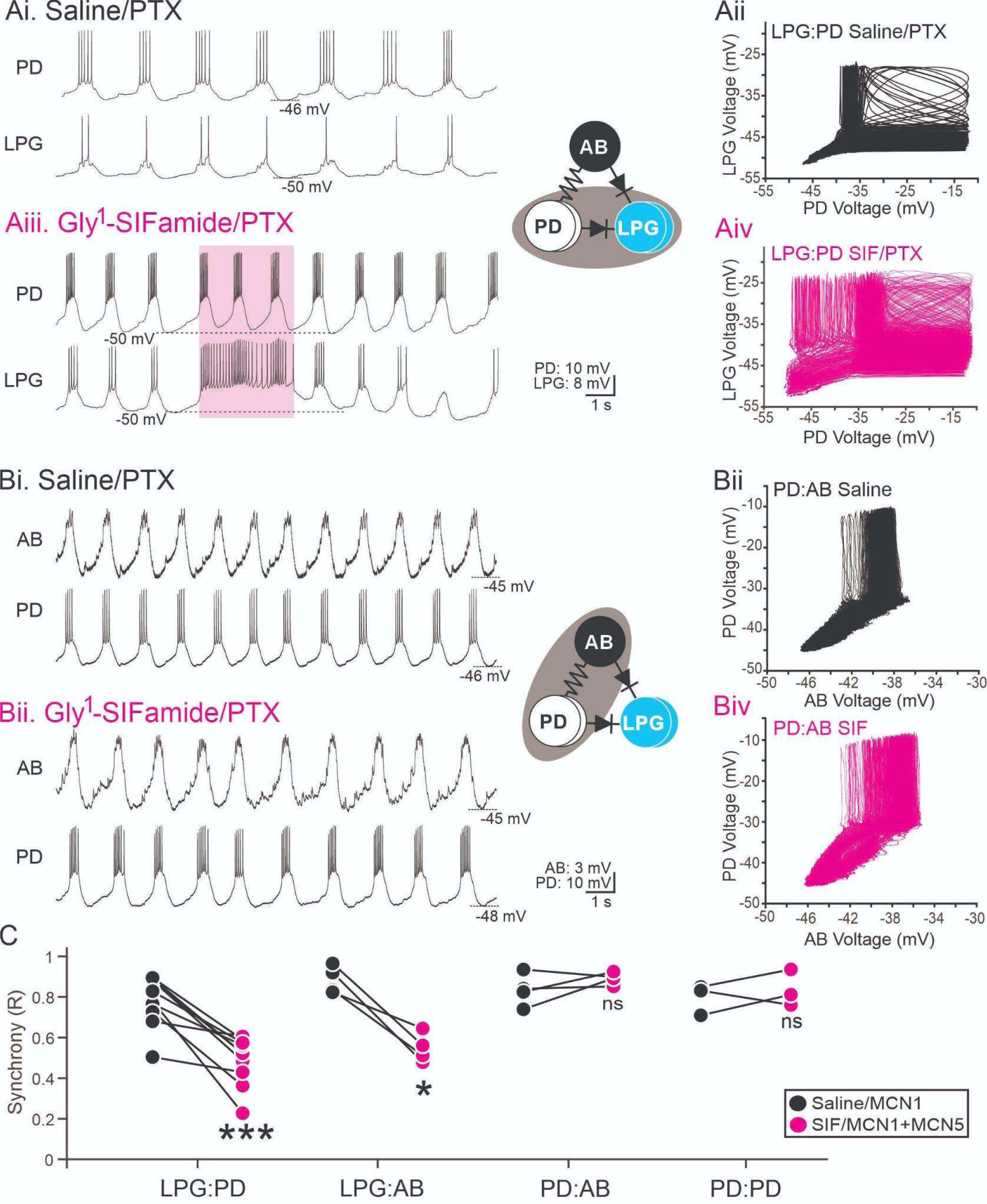
There is a selective decrease in correlated activity between LPG and AB/PD in the MCN5/SIF modulatory state. A) With all pyloric pacemaker neurons intact, a PD and LPG neuron were recorded (grey ellipse) in Saline/PTX (Ai) and during Gly^1^-SIFamide/PTX application (Aiii). Ai) In the control condition, intracellular recordings of PD and LPG in saline plus PTX (10 μM, Saline/PTX) to block glutamatergic input to the pacemaker neurons illustrate similar timing in their slow wave oscillations with different numbers of action potentials per burst. Aii) LPG voltage versus PD voltage is plotted for a 200 s window in Saline/PTX. Aiii) PD and LPG recordings during Gly^1^-SIFamide (5 uM in 10 μM PTX) application in the same preparation as Ai. LPG generated dual network activity that included periodic escapes to generate longer duration gastric mill bursts (example burst indicated by pink box). The voltage trajectory of PD and LPG diverged during this time, with a maintained depolarization in LPG that did not occur in PD (dashes lines indicate trough voltage prior to gastric mill-timed burst). Aiv) LPG vs PD voltage is plotted for 200 s window during Gly^1^-SIFamide/PTX application. Scale bars in Aiii apply to Ai and Aiii. B) With all pyloric pacemaker neurons intact, an AB and PD neuron were recorded (grey ellipse) in Saline/PTX (Bi) and during Gly^1^-SIFamide/PTX application (Biii). AB action potentials are very small in somatic recordings and appear as small bumps at the peak of oscillations. PD voltage was plotted against AB voltage in Saline/PTX (Bii) and in SIF/PTX (Biiii). Recordings in (B) are from a different experiment than those in (A). Scale bars in Biii apply to Biii and Bi. C) The R^2^ values as a measure of synchrony are plotted for 100-200 s windows of LPG:PD (n=10), LPG:AB (n=4), PD:AB (n=4), and PD:PD (n=3) pairs in saline or during MCN1 stimulation (black circles, Saline/MCN1) and during MCN5 stimulation or Gly^1^-SIFamide bath application (pink circles, SIF/MCN1+MCN5). Circles connected by a line indicate data from the same experiment. ***p<0.001, *p<0.05, ns, p>0.05, paired t-test.

Despite the decreased synchrony of LPG and AB/PD neurons, LPG still oscillates in time with AB/PD between its gastric mill bursts, and there appear to be bidirectional influences between AB/PD and LPG, as noted above (e.g., Figs. 1C, 2Aiii) (45), suggesting that electrical coupling between LPG and the other pacemaker neurons is maintained in Gly^1^-SIFamide. However, it was possible that in order to enable LPG to escape, Gly^1^-SIFamide must not only modulate LPG intrinsic currents (28), but it also must decrease the strength of electrical coupling between LPG and AB/PD. Gly^1^-SIFamide does modulate chemical synapses between LPG and gastric mill neurons (46) and electrical coupling can be altered by neuromodulators, including neuropeptides (47, 49, 50). To test whether the Gly^1^-SIFamide neuropeptide modulates electrical coupling among the pyloric pacemaker neurons, we used two-electrode voltage clamp recordings of pairs of LPG and PD neurons.

### Gly^1^-SIFamide does not modulate LPG:PD electrical coupling

To facilitate measurements of electrical synaptic currents, we blocked action potentials with tetrodotoxin (TTX; 1 µM initially, maintained at 0.1 µM) (46).In most modulatory conditions, the pacemaker ensemble only receives chemical synaptic feedback from the LP neuron, however Gly^1^-SIFamide also enhances or enables inhibitory glutamatergic synapses from gastric mill neurons to LPG (46). Thus, LPG and PD were also isolated from LP and gastric mill neuron inhibitory glutamatergic synapses with PTX (10 µM) (8, 33, 51). We use “presynaptic” and “postsynaptic” to distinguish the direction of current flow being examined, even though that designation is arbitrary for bidirectional electrical synapses. We first examined currents elicited in PD (postsynaptic) in response to voltage steps applied to LPG (presynaptic) using two-electrode voltage clamp (TEVC) recordings from one PD and one LPG neuron (Fig. 3Ai). Since these are voltage clamp recordings, the current flow is determined by the difference in voltage between PD and LPG neurons. Thus, the direction of current flow in the postsynaptic neuron is opposite that of the presynaptic voltage step, both above and below - 60 mV where the voltages were the same and there was no net current flow. In an example experiment, in TTX:PTX, a hyperpolarizing voltage step in LPG (-90 mV) relative to PD voltage (-60 mV) elicits an outward (upward) current in PD (Fig. 3Aii, arrows). Further, when PD was held at -60 mV and LPG stepped from -90 to -30 mV, the synaptic currents in PD were larger in the outward (upward) direction, indicating hyperpolarizing current flows preferentially from LPG to PD (Fig. 3Aii, black traces). In the opposite direction, from PD to LPG (Fig. 3Bii), downward (inward) currents were larger in LPG in response to depolarizing voltage steps in PD, than upward (outward) LPG currents in response to hyperpolarizing steps in PD in TTX:PTX, indicating depolarizing current flow preferentially from PD to LPG (Fig. 3Bii, black traces). These results match the previous description of LPG:PD rectification, as well as reports that hyperpolarizing the AB/PD neurons does not alter the LPG trough voltage (8, 35).

**Figure 3.**
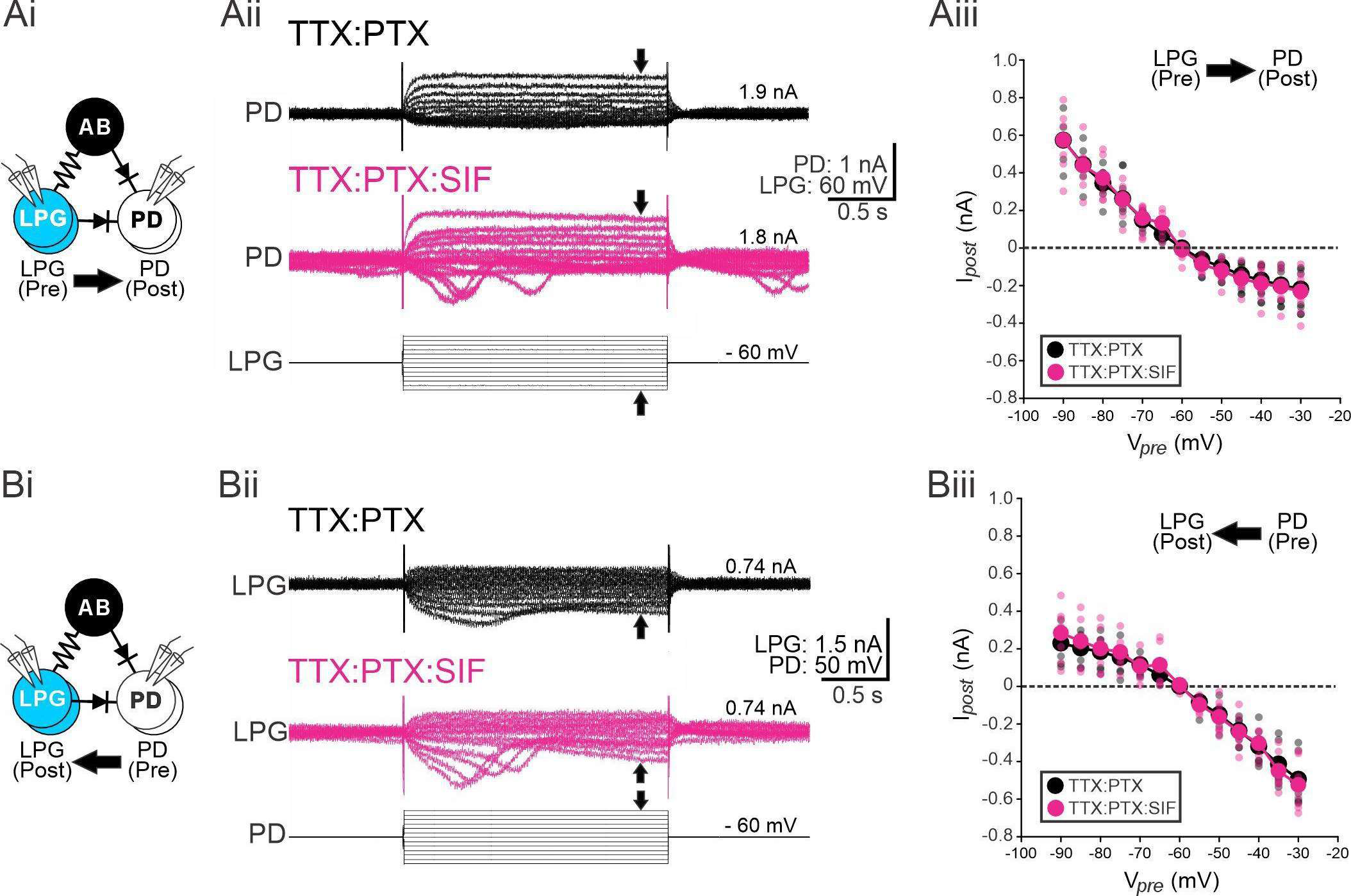
Gly^1^-SIFamide does not alter LPG:PD electrical coupling strength or rectification. Two electrode voltage clamp was performed on a LPG and PD neuron, with other pacemaker neurons unmanipulated. LPG was the “presynaptic” neuron in (A) and PD was the “presynaptic” neuron in (B). Presynaptic refers to the neuron that was stepped across a voltage range and the resulting currents were measured in the “postsynaptic” neuron. Either LPG (Aii) or PD (Bii) was held at -60 mV and stepped from -90 to -30 mV in 5 mV steps and PD (Aii) or LPG (Bii) electrical synaptic currents were recorded in TTX:PTX (black traces) and again in TTX:PTX:SIF (pink traces). The postsynaptic neuron was held at -60 mV between voltage steps. The holding current to maintain all neurons at -60 mV is indicated. The PD (Aiii) or LPG (Biii) electrical synaptic current was measured as the average current during the last 0.8 s of 1 s or the last 1.8 s of a 2 s step minus the baseline current (0.8 s) prior to the voltage step at each voltage level. The average current amplitude across 2-5 trials in each experiment are plotted as smaller semi-transparent dots, and the average across experiments are plotted as larger solid circles (black circles, TTX/PTX; pink circles, TTX:PTX:SIF). There were no differences in the current amplitudes between TTX:PTX and TTX:PTX:SIF. n=7

To test whether the strength or extent of rectification of LPG:PD electrical coupling was different in the MCN5/SIF modulatory state, we bath applied Gly^1^-SIFamide in TTX:PTX (TTX:PTX:SIF) and repeated the voltage protocols. In example recordings of the postsynaptic current in PD or LPG in response to voltage steps in the presynaptic neuron, the strength and rectification of the electrical synapse in TTX:PTX:SIF (Fig. 3Aii, Bii, pink traces) appear similar to those occurring in the control TTX:PTX condition (Fig. 3Aii, Bii, black traces). At the most depolarizing voltage steps during TTX:PTX:SIF application, there were some transient peaks in the inward currents near the beginning of the voltage step in PD (Fig. 3Aii, pink traces) and LPG (Fig. 3Bii, pink traces). These are likely due to depolarizing voltage steps eliciting oscillations in the unclamped PD/AB and LPG. They only occurred in TTX:PTX:SIF and not in TTX:PTX (n=7). In one preparation in which the second PD and the second LPG were photoinactivated, the transient peaks did not occur (data not shown). To quantify the strength and rectification of the electrical synapse, we measured the average amplitude of the synaptic current (the baseline current prior to a voltage step subtracted from the average current during the last 1.8 s of a step) at each voltage. In current-voltage (I/V) plots, the large circles connected by a line indicate the mean current amplitude and smaller semi-transparent circles indicate the individual preparations in TTX:PTX (black) and TTX:PTX:SIF (pink) (Fig. 3Aiii, Biii).

Rectification of the LPG:PD coupling is apparent in the distinct magnitude and slope of the I/V plots above vs below 0 nA. No net current flows between coupled neurons when they are held at the same membrane potential. In our experiments, the postsynaptic neuron was maintained at -60 mV, and thus when the presynaptic neuron was stepped to -60 mV, there was no net current flow between the neurons. To analyze rectification, coupling conductance was determined by fitting the two halves of the I/V relationship (responses to more hyperpolarized [< -60 mV] presynaptic voltages and responses to more depolarized [> -60 mV] presynaptic voltages) and obtaining the slope of each component. From LPG to PD, the coupling conductance was larger for hyperpolarized LPG voltages (Fig. 3Aiii, positive PD currents) (g_elec_ = 18.01 ± 1.51 nS) than for depolarized LPG voltages (Fig. 3Aiii, negative PD currents) (g_elec_ = 8.08 ± 1.45 nS) (n = 7). From PD to LPG, the rectification was in the opposite direction, with a smaller coupling conductance when PD was hyperpolarized relative to LPG (Fig. 3Biii, positive LPG currents) (g_elec_ = 8.54 ± 1.42 nS) than when PD was depolarized relative to LPG (Fig. 3Biii, negative LPG currents) (g_elec_ = 16.28 ± 1.76 nS). Across experiments, there was no change in the magnitude of electrical coupling from LPG to PD (Fig. 3Aii) or from PD to LPG (Fig. 3Bii), as the current amplitudes were similar across voltages between TTX:PTX and TTX:PTX:SIF. Further, there was no change in the coupling conductance in either direction for PD to LPG and thus no change in rectification in TTX:PTX:SIF compared to TTX:PTX (Fig. 3Aiii, Biii). These results indicate that Gly^1^-SIFamide did not modulate electrical coupling strength or rectification between LPG and PD. Therefore, decreased electrical coupling does not explain the ability of LPG to periodically escape from AB/PD oscillations. Instead, we next tested the hypothesis that rectification of the coupling between these neurons is necessary for LPG to escape and generate longer duration gastric mill bursts.

### LPG escapes require rectification of LPG to AB/PD electrical coupling

We first used a mathematical model of the pyloric pacemaker neurons to assess the role of rectification in the LPG escapes. In order to model MCN5/SIF modulation of LPG, we added slow conductances to mimic the Gly1-SIFamide modulated currents (29). With these conductances, when the model LPG is disconnected from AB and PD, it generated slower, longer-duration bursts similar to the biological LPG gastric mill bursts (cycle period: 15.4 s; burst duration 6.6 s) (33) (Fig. 4Ai). LPG was then coupled to the AB and PD neurons with standard symmetrical electrical coupling, and it only oscillated in pyloric time (Fig. 4Aii), despite having the intrinsic conductances to generate gastric mill-timed oscillations when isolated (Fig. 4Ai). However, LPG generated dual-network activity as in the biological network when the AB:LPG, and PD:LPG synapses were instead made rectifying (Fig. 4Aiii). We used coupling conductances based on our measurement of LPG:PD coupling (Fig. 3). These results from the pyloric pacemaker neuron model suggest that electrical synapse rectification is essential for LPG to periodically escape from AB/PD and that non-rectifying coupling prevents the escape. To test this in the biological system, we used the dynamic clamp technique to add simulated symmetrical coupling between PD and LPG.

**Figure 4.**
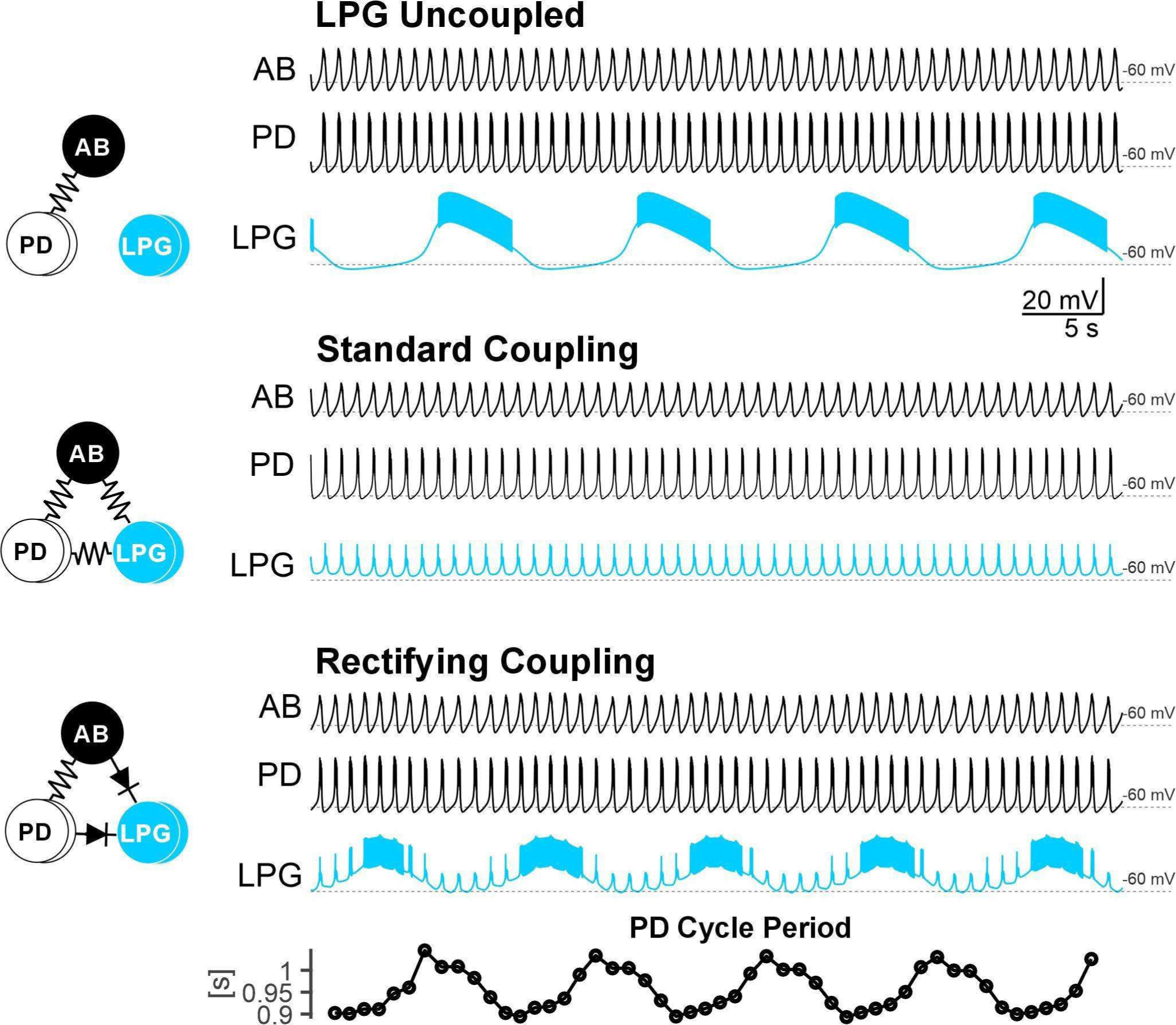
Rectification is necessary for LPG to escape in a computational model of the MCN5/SIF modulatory state. A) Model AB and PD neurons oscillate synchronously due to electrical coupling between them. The uncoupled LPG neuron generates gastric mill-timed oscillations due to simulation of MCN5/SIF modulation (see Methods). B) When the modulated LPG neuron is coupled to AB and PD with non-rectifying electrical synapses, LPG is entirely synchronous with AB and PD (grey box highlights example pyloric-timed oscillation). C) When the AB- and PD-to-LPG electrical synapses in the model are changed to rectifying electrical synapses the LPG neuron produces dual-network activity, with pyloric-timed oscillations synchronous with AB and PD alternating with longer-duration gastric mill-timed oscillations.

To test the impact of a symmetrical LPG:PD electrical synapse on LPG escapes, a single LPG and PD were recorded in two-electrode current clamp to enable accurate voltage measurements separate from current injection. A non-rectifying electrical synapse was modeled in NetClamp software (see *Methods*) in which the amount of current injected into LPG and PD was determined by a coupling conductance (g_elec_) and the differences in their voltages. Addition of a non-rectifying electrical synapses between PD and LPG decreased dual-network activity in LPG such that it remained more synchronous with PD oscillations (Fig. 5, middle). To quantify the impact of adding a non-rectifying electrical synapse, we measured the synchrony (R) in 30 s time blocks before (Pre-Control), during (Dynamic Clamp) and after (Post-Control) a simulated non-rectifying electrical synapse was added between LPG and PD. There was a reversible increase in R with the addition of a non-rectifying electrical synapse (Fig. 5B) (n=3, One way RM ANOVA, Bonferroni post-hoc). In this experiment, we did not alter the existing non-rectifying synapse, however coupling conductances that were only a little larger than the measured coupling conductances in the strong direction were necessary to decrease LPG’s ability to escape (Biological: ∼17 nS, Dynamic clamp conductance: 20-30 nS).

**Figure 5.**
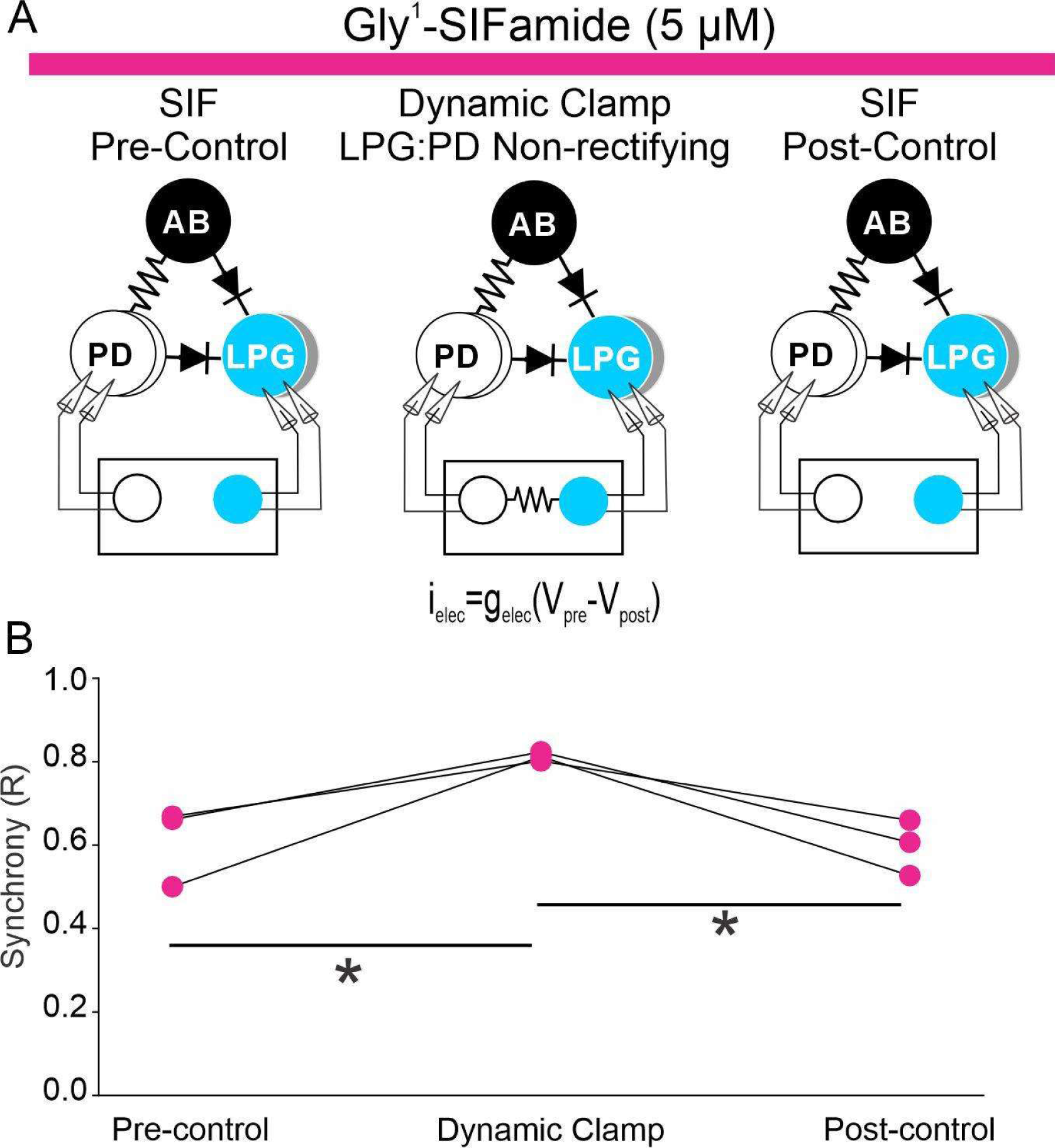
Adding a non-rectifying synapse between LPG and PD decreases the ability of LPG to escape and maintains LPG:PD synchrony. A) In this set of experiments, one PD and one LPG neuron were recorded in two-electrode current clamp mode, with the voltages fed into in a dynamic clamp software (NetClamp, see Methods). A) *Left*, During Gly^1^-SIFamide bath-application, LPG and PD have their typical rectifying electrical synapse. The box represents the dynamic clamp software interface with the two biological cells LPG and PD represented, but not connected. *Middle*, A non-rectifying dynamic clamp electrical synapse (i_elec_=g_elec_[V_pre_-V_post_]; g_elec_=25 nS) was added between LPG and PD via dynamic clamp. LPG and PD voltages are recorded and the appropriate current calculated and injected via the second electrode in each cell. In the “Post-Control” condition, the non-rectifying dynamic clamp electrical synapse was removed B) Plot of PD:LPG synchrony (R) before (Pre-Control), during (Dynamic Clamp) and after (Post-Control) a simulated non-rectifying electrical synapses was added between PD and LPG. *p<0.05, One way RM ANOVA, bonferroni post-hoc.

## Discussion

Here we found that the LPG neuron switching from oscillating at a single frequency to oscillating at two distinct frequencies requires rectification of its electrical synapses to its “home” pyloric network. This was demonstrated in both a computational model and by using a dynamic clamp-simulated non-rectifying electrical synapse in the biological system. In these experiments, the absence of rectification prevented or decreased the ability of LPG to generate oscillations at a second frequency. Instead, LPG maintained synchrony with its home network oscillators, the AB/PD neurons. Additionally, using TEVC analysis of LPG/PD electrical coupling in control versus Gly^1^-SIFamide, we showed that the switch into dual-network activity does not require a decreased strength of LPG electrical coupling to the AB/PD neurons. This means that modulation of LPG’s electrical synapses with its home network, and thus a change in its functional connectivity, is not required to change its network participation. Instead, modulation of the LPG intrinsic properties (8, 28) enables LPG to periodically generate a longer-duration burst and escape from the faster AB/PD neuron bursts, while maintaining the baseline LPG to AB/PD electrical synaptic properties. Thus, the coupling current reaching LPG through the partially rectifying electrical synapses in control conditions is sufficient to synchronize LPG with AB/PD, but not so strong that it prevents LPG from periodically escaping from AB/PD to generate slower, intrinsically-generated bursts in the MCN5/SIF modulatory state.

Maintenance of the strength and rectification of LPG coupling to AB/PD has important consequences for network output and behavior. First, it results in dual-network activity of LPG instead of a complete switch into the slower gastric mill rhythm. LPG innervates both pyloric and gastric mill muscles (31, 52), and at least one of the pyloric muscles can follow LPG dual-network activity (Gnanabharathi, Sheridan, Pinter, Zeigler, Fahoum, and Blitz, unpublished). Second, electrical coupling between LPG and AB/PD enables LPG to contribute to internetwork coordination. Specifically, during the Gly^1^-SIFamide gastric mill rhythm, sustained LPG depolarization during its gastric mill burst increases the pyloric frequency, presumably via the partially-rectified electric synapses from LPG to AB/PD (45). Theoretical studies also supports the role of electrical synapses in coordinating rhythms occurring at distinct frequencies (10). Partial rectification enables LPG to influence AB/PD oscillation frequency, but is sufficiently weak that LPG does not pull AB/PD into a slow burst. A third potential impact of the rectification is that it may play a role in determining LPG sensitivity to other synapses, and thus contribute to network output. In a computational study, replacing a symmetrical synapse with a rectifying electrical synapse greatly constrains network output across a range of strengths of network chemical synapses (15). Gly^1^-SIFamide modulates chemical synapses between LPG and gastric mill neurons, which are necessary for the phase relationships of gastric mill neurons (33, 46), but whether the rectifying electrical synapses regulate the impact of the modulated chemical synapses has not been explored.

It is most intuitive to consider the role of electrical coupling as driving neurons toward synchrony. However, electrical synapses can also be critically important in desynchronizing neuronal activity (25, 53, 54). In fact, electrical coupling can mediate a variety of coordination patterns, in some cases enabling bistability in which an external stimulus can switch coupled neurons between modes of alternation and synchrony (25, 54–56). In the MCN5/SIF state, instead of an external stimulus, neuromodulation alters an intrinsic current(s), to periodically initiate an LPG escape from its electrically coupled partners and generate a slow gastric mill burst. When the burst terminates, LPG returns to its synchronous pyloric-timed activity. Thus, within a single modulatory state the LPG neuron regularly shifts in and out of 1:1 synchrony; this relies on its intrinsic properties (8, 28) and the rectification of its electrical synapses (this study). The combined consequences of electrical synapses and intrinsic properties, as well as chemical synapses, can be quite complex and non-intuitive (14, 16, 25, 57). The potential directionality of electrical synapses due to rectification adds another layer to this complexity (15, 25, 27, 58).

Despite increasing focus on electrical synapses, there remains much to be learned about their contributions to circuit output, including rhythmic CPG networks (14, 16, 25, 59). Although anatomical analyses can provide information about location of electrical synapses, as with chemical synapses, activity-dependent plasticity and neuromodulatory actions, and the potential presence of rectification requires neurophysiological investigation, within the behavioral or modulatory state of interest (14, 53, 57). Here, we add enabling switching between single- and dual-network activity to the litany of roles that electrical synapses play in neural circuits.

## Methods

### Animals and nervous system preparations

The Fresh Lobster company (Gloucester, MA) provided wild-caught adult male *Cancer borealis* crabs. Crabs were housed in tanks containing artificial seawater (10-12 °C) and fed thawed squid twice weekly. Crabs were acclimated at least one week prior to use. Prior to dissection, crabs were anesthetized on ice for 30-50 min. The foregut was dissected out of the animal, bisected, and pinned flat in a Slygard-170-lined dish (Thermo Fisher Scientific) for nervous system dissection. The stomatogastric nervous system (STNS) was dissected free from fat, muscle, and connective tissue and transferred and pinned in a Sylgard 184-lined petri dish (Thermo Fisher Scientific) (8, 60).

All preparations were maintained in chilled (4 °C) *C. borealis* physiological saline throughout dissections, and continuously superfused with chilled *C. borealis* saline (8-10 °C), or chilled saline containing the neuropeptide Gly^1^-SIFamide and/or channel blockers (see Solutions), as indicated. Solution changes were made with a switching manifold to avoid introducing disturbances in solution flow. After obtaining baseline recordings, the *ion* and *son* were cut to remove descending modulatory influences. Preparations were allowed to adapt to the removal of descending inputs for at least 1.5 hr prior to an experiment.

### Solutions

*C. borealis* physiological saline consisted of (in mM): 440 NaCl, 26 MgCl_2_,13 CaCl_2_,11 KCl,10 Trizma base, and 5 Maleic acid (pH 7.4-7.6). A “squid” internal electrode solution was composed of (in mM): 10 MgCl_2_, 400 Potassium D-gluconic acid, 10 HEPES, 15 Na_2_SO_4_, and 20 NaCl (pH 7.45) (61). Aliquots of the neuropeptide Gly^1^-SIFamide (GYRKPPFNG-SIFamide, custom peptide synthesis: Genscript) (12, 28–31) were dissolved in optima water (Thermo Fisher Scientific) at 10^-2^ M. For stock solutions, tetrodotoxin (TTX) was dissolved in optima water (10^-4^ M) and picrotoxin (PTX) (Sigma Aldrich) was dissolved in 95% ethanol (10^-2^ M) and stored at -20 °C. Prior to use, each stock solution was thawed and diluted in physiological saline (final concentration Gly^1^-SIFamide, 5 µM; TTX, 1 µM and 0.1 µM; PTX, 10 µM).

### Electrophysiology

Recordings from peripheral and connecting nerves were made by placing one stainless steel wire from custom-made pin electrodes alongside a nerve, sealing the wire and a small region of nerve with vaseline, and placing the second stainless steel wire outside the sealed region. Differential recordings were obtained with a model 1700 A-M Systems Amplifier. For intracellular recording access to STG somata, the ganglion was desheathed and dark-field (MBL-12010 condenser, Nikon Instruments) illumination used to visualize individual somata. Sharp-tip (18 – 40 MΩ) borosilicate glass microelectrodes were pulled on a Sutter micropipette puller (Flaming/Brown, P-97), and tip- and back-filled with squid internal electrode solution (see *Solutions*) (61) for current clamp recordings, or with 0.6 M K_2_SO_4_ plus 20 mM KCl electrode solution for voltage clamp recordings. Nerve projection patterns, intracellularly-recorded activity and synaptic events, and interactions with other neurons were used to identify STG neurons (2, 62). Intracellular recordings were obtained with AxoClamp 900A amplifiers in current-clamp or two-electrode voltage clamp (TEVC)-mode (Molecular Devices). Current clamp data was recorded with a Micro1401 data acquisition unit (Cambridge Electronic Design) and Spike2 software (∼5 kHz; Cambridge Electronic Design). Voltage clamp data was recorded using a DigiData1440A digitizer (Molecular Devices) and Clampex software (v10.7, Molecular Devices; ∼5 kHz sampling rate). All data was recorded on a laboratory computer (Dell).

In some preparations, MCN5 and/or MCN1 was activated via extraceullular nerve (*ion*) stimulation. MCN1 and MCN5 each project through the *ion* and *stn* to the STG (40, 63). MCN1 typically has a lower voltage threshold than MCN5 and can be selectively activated (64, 65). MCN1 was stimulated at 10-15 Hz to activate a baseline pyloric rhythm and a gastric mill rhythm, which does not include recruitment of LPG into gastric mill activity (39). To activate MCN5, the stimulation voltage was raised to recruit MCN5 and verified through excitatory postsynaptic potentials in AB, inhibition of VD that triggers a strong rebound burst post-stimulation, stronger DG neuron activity, and a switch in the motor pattern, including recruitment of LPG and IC into a gastric mill rhythm (8, 29, 33, 40). The voltage that activated MCN5 also activated MCN1, and thus the projection neurons were coactive.

To mimic MCN5 actions with bath-application of its neuropeptide, we used an established biological model in which MCN5-glutamatergic inhibition of the LP neuron is mimicked by hyperpolarizing or photoinactivating the LP neuron, or blocking the LP inhibitory synapse onto the pyloric pacemaker neurons with picrotoxin (PTX: 10 µM) (8, 28, 33, 45, 46). This mimics MCN5 co-release of Gly^1^-SIFamide, which activates a gastric mill rhythm including modulation of LPG intrinsic properties, and glutamate which inhibits LP (8, 28, 33, 45, 46). For photoinactivation, the LP neuron was impaled with a sharp microelectrode (30-40 MΩ) that was tip-filled with AlexaFluor-568 hydrazide (10 mM in 200 mM KCl, Thermo Fisher Scientific) and back-filled with squid internal solution (see *Solutions*) (8, 33, 45, 66). The LP soma was filled using constant hyperpolarizing current injection (-5 nA) for ∼30 min, or filled for 5-10 min and then the dye was allowed to diffuse to the neurites and axon in the *dvn* for an additional 20-40 min. The STG was then illuminated for 5-7 min using a Texas red filter set (560 ± 40 nm wavelength; Leica Microsystems). Complete photoinactivation was confirmed when the LP membrane potential reached ∼0 mV (8, 33, 45, 66). To test for direct coupling between sets of neurons, or for dynamic clamp experiments (see *Dynamic Clamp*), individual or pairs of pyloric pacemaker neurons were photoinactivated as indicated, using the same method as for the LP neuron.

### Dynamic Clamp

A simulated, non-rectifying electrical synapse was added between a PD:LPG pair of neurons using the dynamic clamp technique (67–69). Dynamic clamp manipulations were performed with NetClamp software (Gotham Scientific) on a Windows 7 PC using a National Instruments PCI-6070-E board. To measure voltage for conductance calculations and also inject current, two-electrode current clamp recordings of a single PD and a single LPG were used. For these experiments, PTX (10 µM) was used to block chemical synaptic input to the PD and LPG neurons. Additionally, in some dynamic clamp experiments the second LPG neuron was photoinactivated to enable all LPG activity to be directly influenced by the dynamic clamp current addition. The second PD neuron and AB were always intact. A bidirectional non-rectifying electrical synapse was modeled with the equation: i_elec_=g_elec_(V_pre_-V_post_). As the synapse was bidirectional, this equation was used to inject current into the PD neuron with LPG as the presynaptic and PD as the postsynaptic neuron, and into the LPG neuron with PD as the presynaptic neuron and LPG as the postsynaptic neuron. A maximal conductance (g_elec_) that was effective ranged from 10-30 nS. This range overlaps with the conductance values obtained for the PD:LPG electrical coupling in this study.

### Data Analysis

To assess neuronal synchrony, the voltages of pyloric pacemakers were recorded in control saline or during *ion* (MCN1 only) stimulation and compared to their voltage trajectories during Gly^1^-SIFamide (5 µM) bath application or *ion* stimulation (MCN1 + MCN5). In all conditions, neuronal activity was recorded unperturbed for 100-200 s after reaching a steady state level. As a measure of synchrony, the correlation coefficient of the voltages of neuron pairs (LPG:AB, LPG:PD, AB:PD, PD:PD) was calculated using a custom MATLAB (r2024a; MathWorks) script (47, 48). This was performed for control conditions and during addition of dynamic clamp currents.

Electrical coupling currents were measured by holding the postsynaptic cell at -60 mV and stepping the presynaptic cell from -90 to -30 in 5 mV steps for 2 s. Cells were held at -60 mV between runs. The average current across 1 s prior to the step was subtracted from the average current across the last 1.8 s of each voltage step. Averages of 2-3 trials per experiment were averaged across experiments. Coupling conductances were calculated by fitting each half of the V/I curves (below and above -60 mV holding potentials).

### Software and statistical analysis

Graphs were made in Matlab (r2024a; Mathworks) or SigmaPlot (v14; Systat Software) and imported into CorelDraw (v24; Corel Corporation) where all final figures were made. Statistical analysis was performed with Sigmaplot. Data were first assessed for normality using the Shapiro-Wilk test, followed by paired t-test or one-way repeated measures ANOVA as indicated. An alpha level of 0.05 was used as threshold for statistical significance. Data reported as mean ± standard error.

### Code accessibility

Scripts used for analysis are available at https://github.com/blitzdm/FahoumNadimBlitz.

## Acknowledgements

NSF IOS 1755283 (DMB)

## Supporting Information

**Figure S1.**
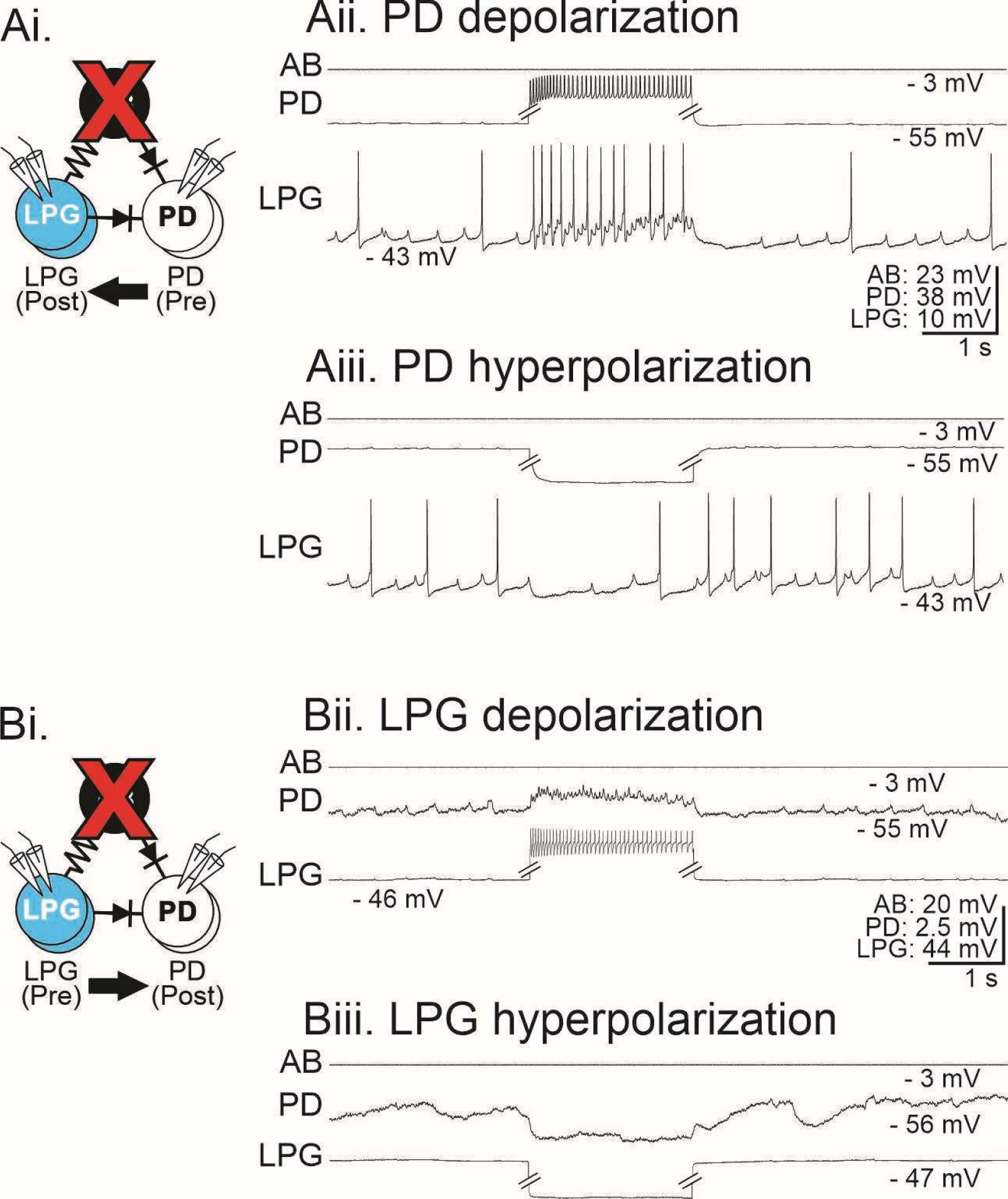
LPG and PD were directly electrically coupled in the absence of AB. The AB neuron was photoinactivated (Ai, Bi; red X) and depolarizing and hyperpolarizing current injected into PD (Aii-Aiii) or LPG (Bii-Biii) to test for electrical coupling. Current injections were performed with an explicitly unbalanced bridge circuit and traces were truncated for ease of visualization. Double lines indicate breaks in voltage traces in PD (Aii-Aiii) and LPG (Bii-Biii).

